# Septins and a formin have distinct functions in anaphase chiral cortical rotation in the C. *elegans* zygote

**DOI:** 10.1101/2020.08.07.242123

**Authors:** Adhham Zaatri, Jenna A. Perry, Amy Shaub Maddox

## Abstract

Many cells and tissues exhibit chirality that stems from the chirality of proteins and polymers. In the *C. elegans* zygote actomyosin contractility drives chiral rotation of the entire cortex circumferentially around the division plane during anaphase. How contractility is translated to cell-scale chirality, and what dictates handedness, are unknown. Septins are candidate contributors to cell-scale chirality because they anchor and crosslink the actomyosin cytoskeleton. We report that septins are required for anaphase cortical rotation. In contrast, the formin CYK-1, which we found to be enriched in the posterior in early anaphase, is not required for cortical rotation, but contributes to its chirality. Simultaneous loss of septin and CYK-1 function led to abnormal and often reversed cortical rotation. Our results suggest that anaphase contractility leads to chiral rotation by releasing torsional stress generated during formin-based polymerization, which is polarized along the cell anterior-posterior axis, and which accumulates due to actomyosin network connectivity. Our findings shed light on the molecular and physical bases for cellular chirality in the *C. elegans* zygote. We also identify conditions in which chiral rotation fails but animals are developmentally viable, opening avenues for future work on the relationship between early embryonic cellular chirality and animal body plan.

## Introduction

Chirality (or “handedness”) is an intrinsic feature of proteins, subcellular structures, cells, organs and organisms. Large scale chirality can emerge from fluid flows and contraction (reviewed by (Pohl, 2015). Cellular chirality is attributed to the chirality of constituent proteins and other biomolecules, but how this asymmetry or handedness is translated several orders of magnitude in length scale from molecule to cell is poorly understood. The polymerization of cytoskeletal proteins into much larger filaments is an excellent candidate for the scaling-up of chirality.

*C. elegans* undergoes deterministic development; blastomeres often adopt specific fates upon their formation during cleavages and body axes are established early (Sulston et al., 1983). Even in the one-cell embryo (zygote), where only the anterior-posterior axis is initially apparent, several rotational cortical flows with consistent chirality occur. In early anaphase, the entire cortex of the cell, associated cortical granules, endoplasmic reticulum, and microtubules, rotates (Pimpale et al., 2020; Schonegg et al., 2014; Singh et al., 2019; Singh and Pohl, 2014). The cortex flows around the anterior-posterior axis with a right-handed chirality; particle movement resembles the curl of right-hand fingers when the right hand thumb is pointed to the posterior (Figure 1A) (Naganathan et al., 2014; Schonegg et al., 2014). This cortical rotation depends on zygote anterior-posterior polarity and is implicated in the cortical chirality of early blastomeres and establishment of the dorso-ventral body axis, but is independent of the left-right axis of the body plan (Schonegg et al., 2014). While other asymmetrically-dividing early blastomeres also exhibit this “net-rotating flow” in which the entire cortex rotates in the same direction, symmetrically-dividing blastomeres instead exhibit chiral counter-rotation, during which cortex in the posterior half flows around the circumference with right-handed chirality, while cortex in the anterior half flows with left-handed chirality (Pimpale et al., 2020). During embryo polarization, cortex flows not only from the posterior to the anterior (Hird and White, 1993), but also in a counter-rotating fashion, with the posterior half exhibiting right-handedness and the anterior half flowing with a left-handed chirality (Naganathan et al., 2014). Finally, during cytokinesis, the cortex flows into the cell equator, as observed throughout phylogeny (Khaliullin et al., 2018; White and Borisy, 1983). Cortical flows affect spindle positioning and therefore division plane specification and the placement and intercellular contacts of the resulting cells (Sugioka and Bowerman, 2018). Thus, defining the mechanisms controlling and executing flows may generate insights into developmental morphogenesis.

**Figure 1:**
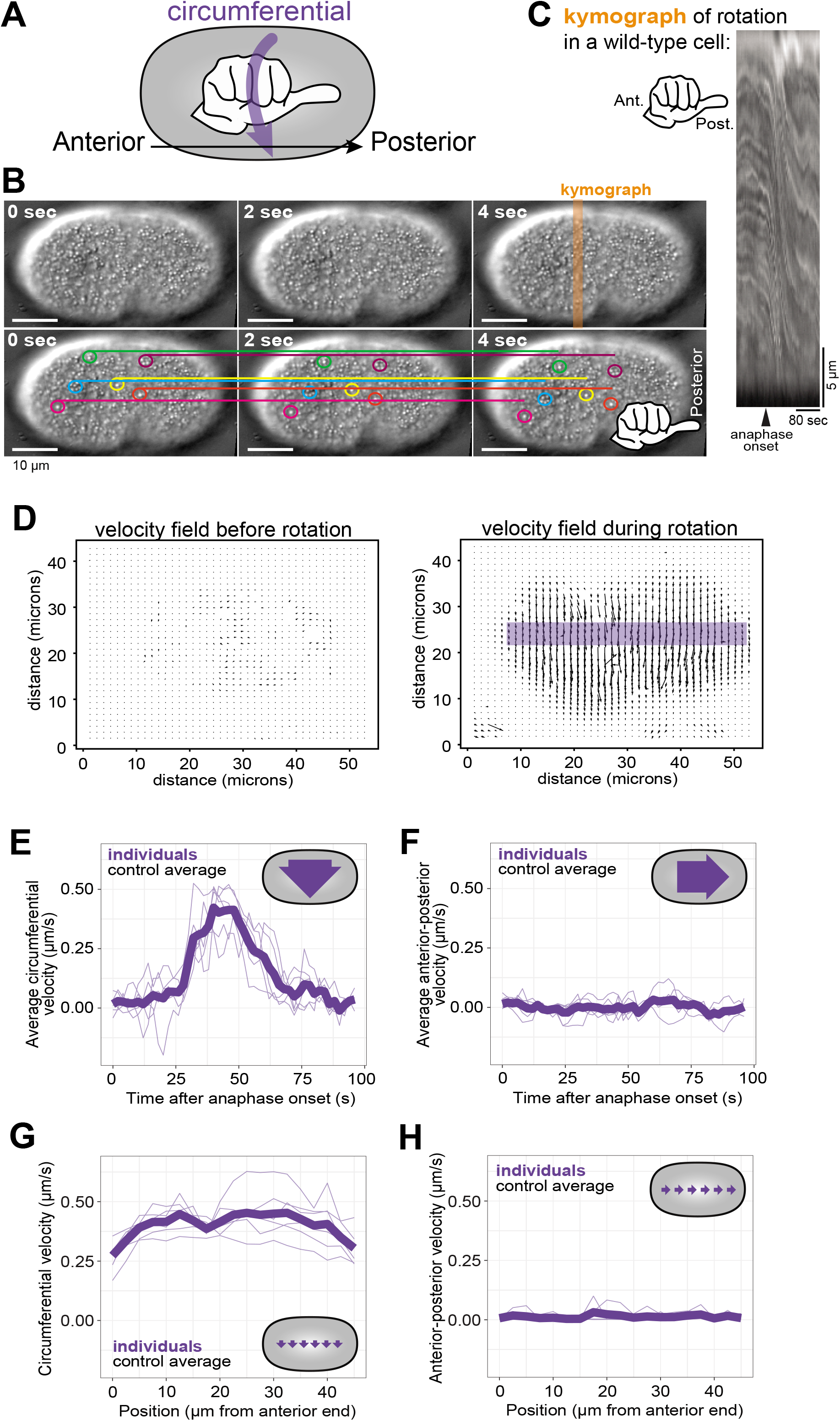
Cortical rotation in *C. elegans*. (A) A schematic depiction of the use of a hand with thumb pointed towards the zygote posterior to determine the chirality of *C. elegans* cortical rotation. (B) Consecutive frames of cortical rotation (dt = 2s). Colored circles highlight cortical granules, while the corresponding colored line denotes the starting location (at t = 0s) of the granule across all time points. Scale bar = 10 microns (C) Kymograph of cortical rotation in a control cell. (D) Representative vector field outputs from OpenPIV using the same cell as in (A), before and during rotation. Arrow length represents local velocity. (E-F) Circumferential (around the anterior-posterior axis) and Posterior-directed velocities during cortical rotation over time, averaged across the length of the cell. (G-H) Averaged circumferential velocity of cortical rotation with respect to the position along anterior-posterior axis. Pale purple lines are individuals; heavy purple line is the population average (n = 6).

Cortical rotation in the anaphase *C. elegans* zygote depends on the actomyosin and microtubule cytoskeletons (Schonegg et al., 2014). Non-muscle myosin II drives contractility and remodeling of the actin cytoskeleton (Koenderink and Paluch, 2018). The activity levels and spatial patterning of both actin polymerization and non-muscle myosin II are controlled by the small GTPase RhoA, which itself is patterned by the microtubule cytoskeleton (Koenderink and Paluch, 2018). Further mechanistic insight into cellular chirality is gained from work with adherent cultured mammalian cells in which radial actomyosin bundles form upon cell spreading. These initially-radial bundles gradually adopt a chiral tilt, in a manner dependent on actomyosin contractility and actin polymerization by formins, which act in association with the membrane where they are activated by RhoA (Tee et al., 2015). This cell chirality also depends on crosslinking of radial and circumferential cytoskeletal bundles.

Septins are cytoskeletal proteins that form membrane-associated polymers that directly bind several components of the cortical cytoskeleton including F-actin and non-muscle myosin II. Septins have roles in cell division and cytoskeletal remodeling (Weirich et al., 2008) via their contributions to the cortical localization of many cytoskeletal components and their regulators (Joo et al., 2007; Spiliotis and Gladfelter, 2012). In this capacity, septins contribute to cellular asymmetries (Gilden and Krummel, 2010; Maddox et al., 2007; Mostowy and Cossart, 2012) and to symmetry-breaking of actin-based structures (Maddox et al., 2007; Mavrakis et al., 2014; Spiliotis, 2010). *C. elegans* septins enrich in the zygote anterior but are dispensable for zygote anterior-posterior polarity (Davies et al., 2016; Jordan et al., 2016; Nguyen et al., 2000). They enrich in the cytokinetic ring but are also generally dispensable for cytokinesis in *C. elegans*, though septin depletion causes the zygote cytokinetic ring to close concentrically, not unilaterally, as in control cells (Maddox et al., 2007; Nguyen et al., 2000). The *C. elegans* genome has only two septin genes, *unc-59* and *unc-61*, which encode proteins that form a non-polar heterotetramer composed of UNC-59 at the ends and UNC-61 in the middle (John et al., 2007; Nishihama et al., 2011). Septins are candidates for crosslinking actin-based structures and for scaffolding formin-generated F-actin at the membrane (Akhmetova et al., 2018; Breitsprecher and Goode, 2013; Buttery et al., 2012; Gao et al., 2010). The role of septins in cortical rotation has been examined in highly compressed *C. elegans* zygotes that are sensitized to fail cytokinesis, and in which the integrity of longitudinal (anterior-posterior) myosin bundles is implicated in rotation (Singh et al., 2019). Septins are required for rotation in this condition, but their role in cellular chirality in less deformed cells remains unknown (Singh et al., 2019).

Here we compared wild-type *C. elegans* zygotes with those depleted of the septin UNC-59, and from strains with loss-of-function mutations in both genes that encode septins, *unc-59* and *unc-61*. Qualitative assessment of cortical rotation as well as quantification by particle image velocimetry (PIV) (Liberzon, 2019) demonstrated that septins are required for cortical rotation. To understand how septins transmitted polymer or molecular chirality to the cellular level, we interrogated the interplay between septins and the formin CYK-1. We observed an enrichment of GFP-tagged CYK-1 in the zygote posterior that was lost when septins were depleted. Loss of CYK-1 function also perturbed cortical rotation, but in a manner distinct from the effects resulting from septin loss of function. Simultaneous loss of function of a septin and CYK-1 largely eliminated rotation, and caused randomization of rotation chirality. These findings begin to elucidate the roles of septins and CYK-1 in driving and controlling chiral rotation and suggest that septin scaffolding of the actomyosin cytoskeleton contributes to cellular chirality.

## Methods

### *C. elegans* strains and maintenance

*C. elegans* (see Table 1 for strain names and genotypes) were maintained on nematode growth medium (NGM) and OP50 bacterial food at 20°C. Worms were transferred to new plates two days prior to imaging in non-RNAi experiments.

**Table 1:** Genotypes of *C. elegans* strains

| Strain | Genotype |
| --- | --- |
| N2 | Wild type |
| JCC239 | *unc-61(e228)* V; *unc-119(ed3)* III; ltIs37 [pAA64; *pie-1*/mCHERRY::his-58; *unc-119* +] IV |
| JCC661 | *unc-59(e1005)* I.; *unc-119(ed3)* III*; ltIs38 [pAA1; *pie-1*/GFP::PH(PLCdelta1); *unc-119* (+)] III.; *unc-119* (+) III; ltIS37 [pAA64; *pie-1*/mCherry::his-58; *unc-119*(+)] IV. |
| MDX79 | ltIs37 [pAA64; *pie-1* /mCHERRY::his-58; *unc-119* (+)] IV; *cyk-1(ges1[cyk-1::eGFP+LoxPunc-119(+)]LoxP)* III; *unc-119(ed3)* * III. |
| MDX82 | SWG001 x JCC719 mgSi3[*cb-unc-119* (+); *pie-1::gfp::utrophin*] II; gesls0001[*Mex5p::lifeact::mKate*; *unc-119* (+)]. |
| NK2228 | *unc-59(qy88[unc-59::GFP::3xflag::AID+loxP])* |

\* since this lesion is rescued by transgenes, its maintained presence is unknown

The strains expressing CYK-1::GFP and UNC-59::GFP were previously published (Chen et al., 2019; Hirani et al., 2019). The strain bearing *unc-59(e1005)*, characterized and kindly provided by Julie Canman, bears G160-to-A160, not G85-to-A85 (encoding UNC-59 with Gly54-to-Arg54 not Gly29-to-Arg29) as previously described (Nguyen et al., 2000).

### RNA-mediated protein depletion (RNAi)

Starved worms were grown on NGM-OP50 for 24 hours at 20°C. 10-20 fourth larval stage (L4) worms were transferred onto an NGM plate containing 1mM IPTG and 0.132 mM carbenicillin, and seeded with bacteria expressing dsRNAs targeting *unc-59* and *cyk-1*. Imaging of dissected zygotes began 20 hours from the start of RNAi feeding for CYK-1 depletion, or 24 hours for UNC-59 depletion.

### Embryo mounting and image acquisition

Worms were mounted onto 2% agarose pads as previously described except that they were transferred from a drop of M9 solution on a cover glass and not via mouth pipet (Maddox and Maddox, 2012).

The same procedure was employed in mounting zygotes for measuring intensity of CYK-1::GFP except for the photobleaching control cells that were instead dissected in 1.5 μL of 0.1 M of Na++-Azide in M9 buffer.

For Figures 1-3, a DeltaVision Elite microscope (Cytiva) with an Olympus 60X 1.40 NA silicone oil immersion objective lens was used to image embryos by differential interference contrast. Every 2 seconds, two optical sections were acquired: the midplane of the cell (for timing anaphase onset) and the cortical plane proximal to the coverslip. For Figure 3, CYK-1::GFP localization before rotation was imaged on the DeltaVision, and CYK-1::GFP timelapse image series with a 3-second temporal interval were acquired on a Nikon A1R scanning confocal using a 60X 1.27 NA water immersion objective lens. For any given figure panel, identical acquisition settings were used for all conditions.

**Figure 3:**
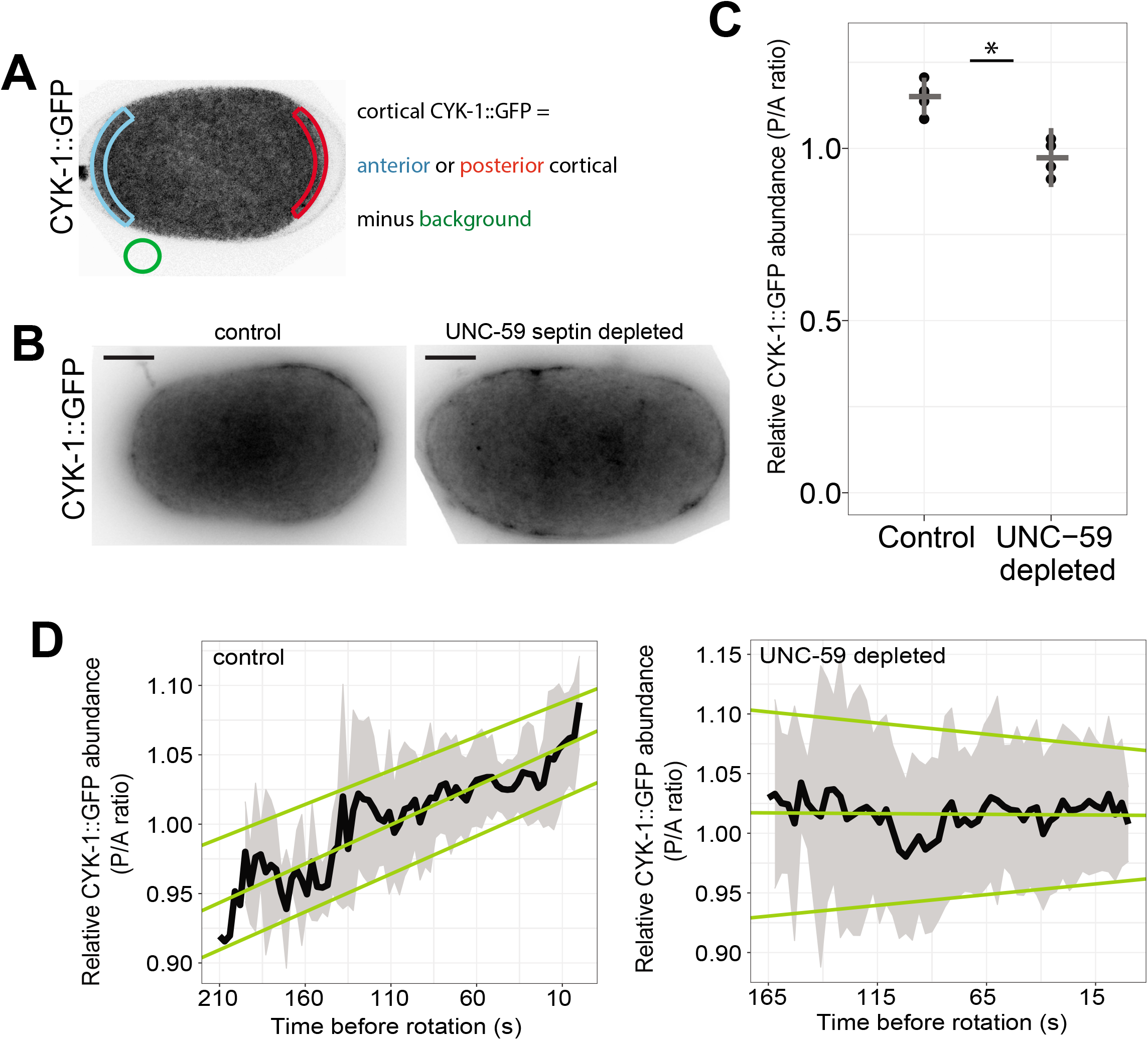
Formins have a septin-dependent posterior-enrichment in the *C. elegans* zygote. (A) Schematic of intensity measurements. (B) Control and *unc-59* depleted cells exhibited CYK-1::GFP. Scale bar = 10 microns. (C) Mean intensity ratio (P/A) of CYK-1::GFP just prior to anaphase onset for control (n = 5) and *unc-59(RNAi)* (n = 4) cells. Dots are individuals; horizontal bar is population mean; vertical notches are 95% confidence interval. (*) p < 0.01. (D) Mean intensity ratio of CYK-1::GFP (P/A) leading up to cortical rotation. Black line is population mean (control: n = 12, UNC-59 depleted: n = 5). Gray shaded areas are 95% confidence intervals; pale green lines are linear regression for upper, middle, and lower bounds of confidence.

To determine the handedness of cortical rotation imaged on our DeltaVision Elite microscope, we used the orientation of a printed letter “F” mounted on a microscope slide as a guide. In raw images, the letter F appeared mirrored, as compared to when viewed with the naked eye. To generate a true-to-life image orientation, therefore, we mirrored our images of *C. elegans* zygotes (in which cortical rotation had appeared to have left-handed chirality).

### Image Analysis

For each time series, the cortical plane sections were separated into a series of TIFF images using Fiji. The series of cortical images was then analyzed using OpenPIV, a Particle Image Velocimetry (PIV; http://www.openpiv.net/) plugin for Python (Liberzon, 2019; Schindelin et al., 2012). From the resulting vector field of the velocities, vectors perpendicular and parallel to the cell long axis were separately averaged.

The frames during which rotation occurred were designated as those when circumferential velocity was at least 0.09 micron/s, with a tolerance 0.01 micron/sec lower for the frame immediately before/after the initially determined rotation period. Circumferential velocity is defined as speed in the right-handed direction perpendicular to the anterior-posterior axis. Anterior-posterior velocity is defined as speed directed towards the zygote posterior. Time frames within the tolerance were discarded if the velocity cutoff criterion was met for less than eight seconds, since in those cases the heightened speed could likely be attributed to stochastic movement or artifacts.

Average velocity was measured with a region 5 microns tall centered a manually defined central line of the cell (see Figure 1D, purple bar). Positional velocity measurements averaged the velocity over the whole rotation interval by dividing the aforementioned region of interest into 20 bins along the anterior-posterior axis. The anterior-posterior axis was identified via the presence of polar bodies (anterior) and displacement of the anaphase spindle (posterior). Rotation type (as defined below) was determined using the average velocity measurements. Continuous rotation had a single period of at least 38 seconds of circumferential velocity above 0.09 microns/sec. Intermittent rotation had at least one period, lasting at least 8 seconds, above the aforementioned velocity cutoff, as well as at least one period of non-negative velocity below the cutoff. Alternating rotation had at least one period of rotation with circumferential velocity above 0.09 microns/sec followed by a period with a velocity of −0.09 microns/sec or below (in the opposite direction of rotation), each at least 8 seconds long. No rotation meant that no frames met the previously described criteria for defining a period of rotation.

Fluorescence intensity for time series acquired on the DeltaVision Elite was measured using Fiji. Background fluorescence intensity outside the cell was subtracted from all intensity measurements in the cell (see Figure 3A). For images recorded on the A1R, intensity was measured by specifying regions of interest pre-imaging and exported directly from the NIS-Elements imaging software (Nikon Instruments). The change in intensity over time was fitted to an exponential decay curve.

Kymographs were made using Fiji’s multi kymograph tool. A 10 pixel-wide line was drawn perpendicular to the anterior-posterior axis such that the line originated from the top of the image, with anterior on the left.

### Statistical Analysis

Pairwise Student’s t-tests were performed in opensource R software for all comparisons of perturbations with the corresponding controls, except for data presented in Fig. 3 supplement A and Fig. 4I, which were analyzed via t-test and ANOVA using Prism software.

**Figure 4:**
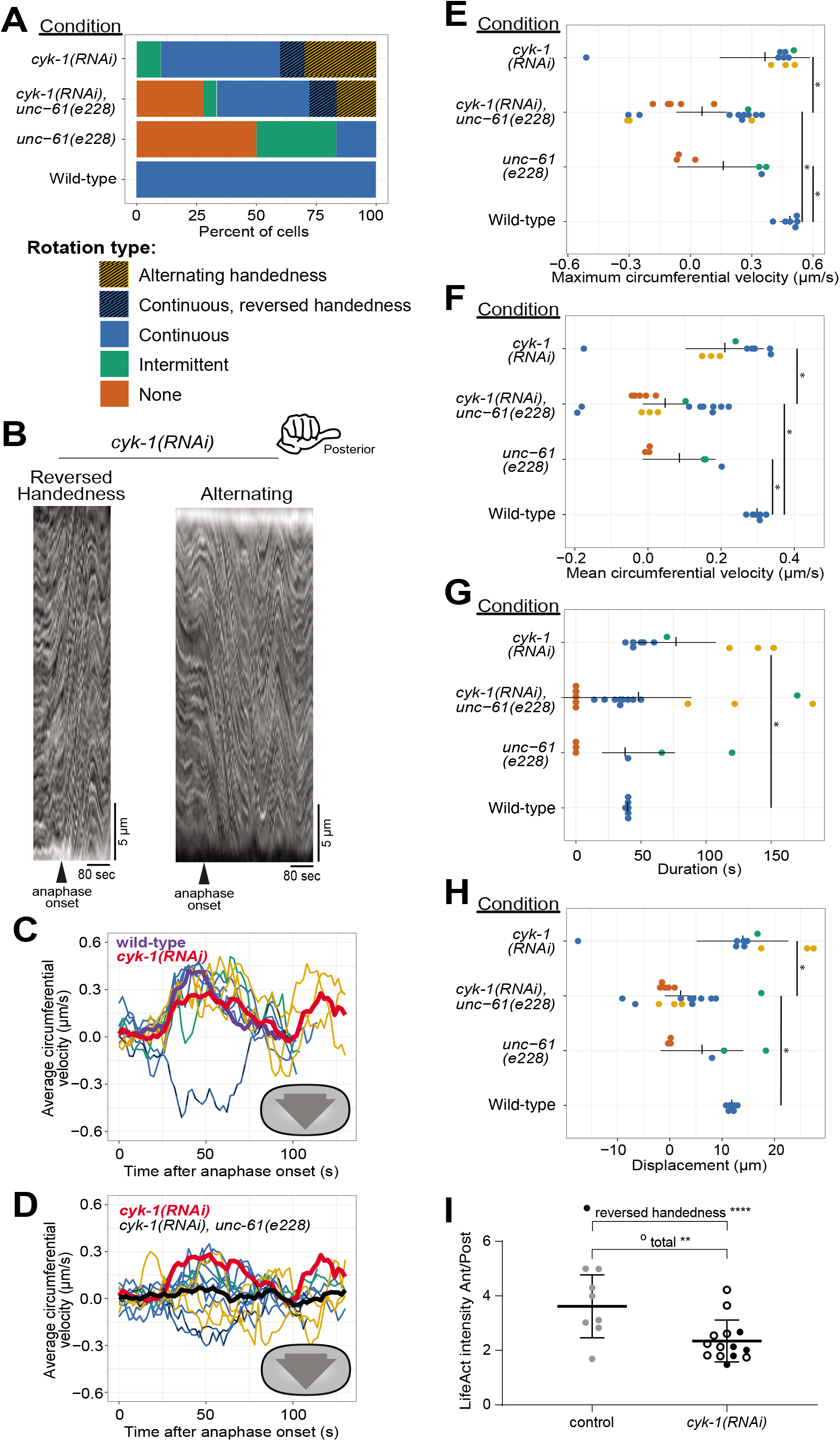
Cortical rotation is dependent on the formin CYK-1. (A) Phenotype frequency for various genetic perturbations of *unc-61* and *cyk-1*. Alternating rotation changes direction at least once; continuous rotation has a single period of increased velocity; intermittent rotation has several periods interspersed with periods of slower or no movement; none indicates that no rotation occurred. (B) Example kymographs of cells exhibiting reversed handedness or alternating rotation. (C-D) Circumferential velocities over time averaged across the anterior-posterior axis. Colored lines indicate individuals of the corresponding color-coded phenotype; purple: control average; red: *cyk-1(RNAi)* population average (n = 10); brown: (*cyk-1(RNAi); unc-61(e228))* population average (n = 18); dashed line: reversed handedness. (E-H) Quantification of mean velocity, maximum velocity, duration, and displacement, respectively. Colored dots are individuals of the corresponding phenotype. Vertical notches are population means; horizontal lines are 95% confidence intervals. (*) = p < 0.05; unmarked pairings are not significantly different. (I) Ratio of LifeAct fluorescence in the zygote anterior versus posterior in anaphase in control (grey) and CYK-1 depleted cells (black; open: continuous and intermittent rotation with right handed chirality, filled: rotation with continuous left-handed chirality (n=3) or alternating handedness (n=2)) (**) p < 0.01; (****) p < 0.001.

## Results

### Quantifying cortical rotation

Cortical rotation in the *C. elegans* zygote is an example of cell-scale chirality whose mechanism may provide insights into how molecular chirality influences cell behavior. To uncover molecular mechanisms of this event, we first quantitatively analyzed zygote anaphase cortical rotation. We began by using DIC transmitted light microscopy to visualize zygotes during late metaphase and early anaphase. Cytoplasmic granules near the cortex of control *C. elegans* zygotes were displaced circumferentially around the cell, their trajectories exhibiting a right-handed chirality around the anterior-posterior axis (Figure 1A-C; Supplemental Movie 1), consistent with published findings (Schonegg et al., 2014).

We used particle image velocimetry (PIV) to quantify cortical rotation by tracking the motion of particles between pairs of images, identifying patches of texture, such as cytoplasmic granules (Figure 1B), in the first image, then finding those same patches in the second image. PIV then generates a vector field of the velocity of features in the images based on their displacement and the time between frames (Figure 1D). The vectors, representing the motion of cortical granules, pointed around the cell circumference (around the anterior-posterior axis) with right-handed chirality.

The average velocity during 25 seconds of cortical rotation has been reported as 0.35 microns/second (Singh and Pohl, 2014). However, since particles are essentially stationary before anaphase cortical rotation, and rotation has a finite duration of 50-60 seconds (Schonegg et al., 2014), we sought to characterize the speed dynamics of rotation throughout its duration. We used the spatially-and temporally-resolved data from the vector field and found that cortical rotation first accelerated and then decelerated, peaking near 0.4 microns/second, approximately 40 seconds after anaphase onset (Figure 1E). For any given timepoint during rotation, the vector field was notably uniform, with only slight variations in angle relative to the anterior-posterior axis. The average instantaneous velocity in the anterior-posterior direction was very low (Figure 1F). As previously described (Schonegg et al., 2014; Singh and Pohl, 2014), rotation velocity was relatively uniform along the length of the embryo (Figure 1G, H). Thus, the entire *C. elegans* zygote cortex undergoes a short-lived chiral rotation that displaces cortical material around the cell circumference during anaphase.

### Septins are required for cortical rotation

Cortical rotation in the *C. elegans* zygote is driven by contractility of the actomyosin cytoskeleton associated with the plasma membrane (Schonegg et al., 2014). Septins are implicated in linking the cytoskeleton to the membrane. Septins also contribute to cell cortex polarity in yeast (Bridges and Gladfelter, 2015) and furrowing asymmetry in the *C. elegans* zygote (Maddox et al., 2007). In the *C. elegans* zygote, septins are enriched in the anterior cortex and in the cytokinetic ring (Jordan et al., 2016; Nguyen et al., 2000) (Figure 3 supplement, A). *C. elegans* septins are required for zygote anaphase chiral rotation in severely compressed cells, but are not required for the chiral counter-rotation during zygote polarization (Naganathan et al., 2018; Singh et al., 2019). Since cortical mechanics are affected by cell compression and therefore shape (Singh et al., 2019), we tested whether septins were required for zygote anaphase cortical rotation in cells that were not deformed beyond the slight compression arising from traditional mounting of *C. elegans* embryos (Pimpale et al., 2020).

We first depleted the septin UNC-59 via RNA-mediated interference (RNAi) and found that in many UNC-59-depleted cells, cortical rotation failed (Figure 2A, C, F& G, Figure 2 supplement, and Supplemental Movie 1). In UNC-59-depleted zygotes undergoing anaphase cortical rotation, both the maximal and average rotation velocities were reduced (Figure 2C, F, G). The duration of rotation was reduced in most cases (Figure 2H) and total distance was significantly reduced compared to controls (Figure 2I). Thus, as in severely compressed cells that fail cytokinesis (Singh et al., 2019), septins are required for normal cortical rotation in cells that complete cytokinesis (Maddox et al., 2007).

**Figure 2:**
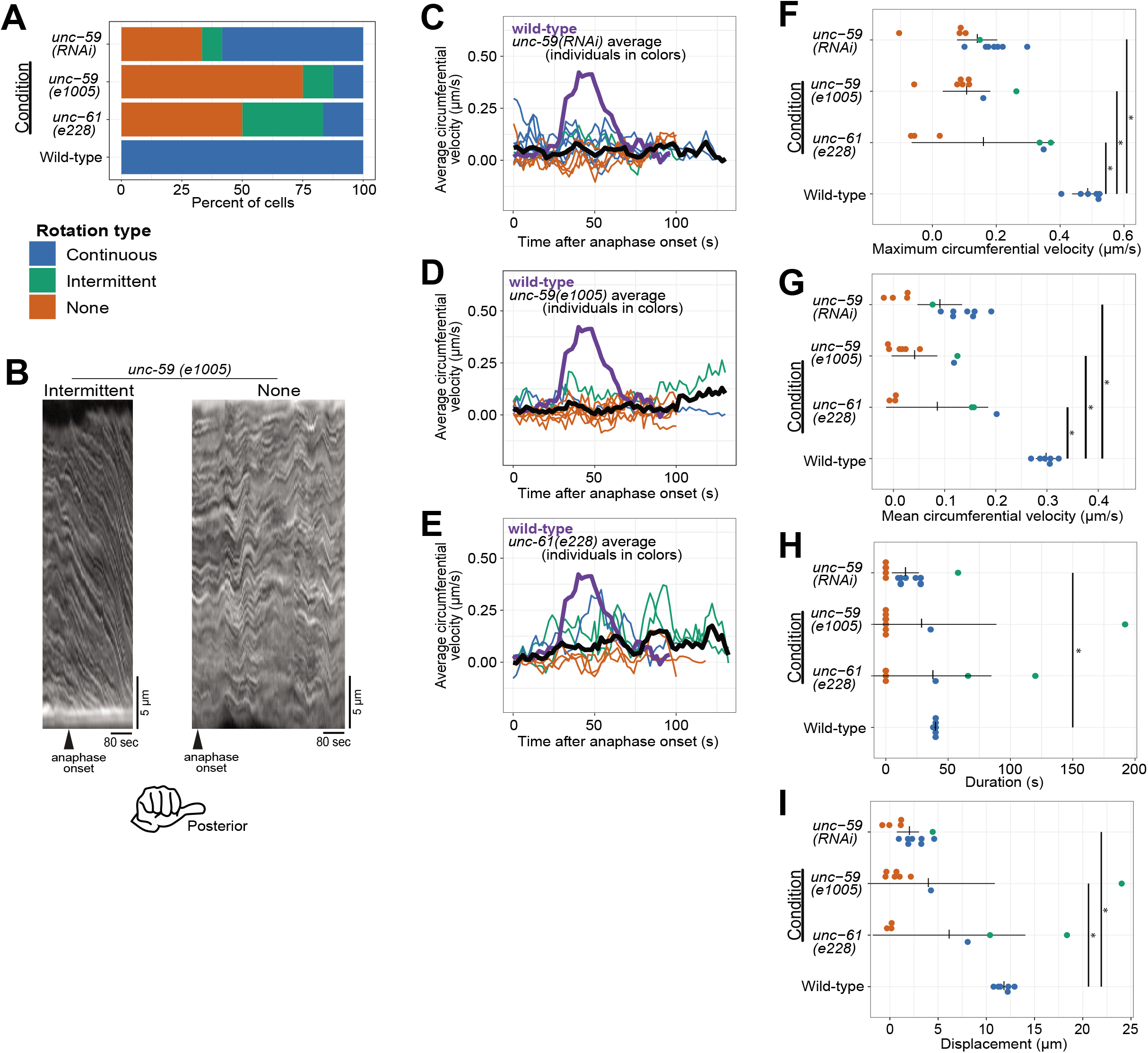
Cortical rotation is dependent on septins. (A) Rotation phenotype frequency for genetic perturbations of septins. Continuous rotation has a single period of increased velocity; intermittent rotation has several periods interspersed with periods of slower or no movement; none indicates that no rotation occurred. (B) Example kymographs of cells exhibiting intermittent or no rotation (see Figure 1 for continuous). (C-E) Circumferential velocities over time averaged across the anterior-posterior axis. Colored lines indicate individuals of the corresponding color-coded phenotype; black line is the population average (*unc-59(RNAi)* n = 12; *unc-59(e1005)* n = 9; *unc-61(e228)* n = 6); purple line is the control average (n = 6). (F-I) Quantification of mean velocity, maximum velocity, duration, and displacement, respectively. Colored dots are individuals of the corresponding phenotype. Vertical notches are population means; horizontal lines are 95% confidence intervals. (*) p < 0.05; unmarked pairings are not significantly different.

We next tested the effect of septin loss of function using *C. elegans* zygotes bearing a mutant allele of *unc-59*. This allele encodes a point-mutated UNC-59 protein that fails to localize to the cytokinetic ring (*e1005*; (Nguyen et al., 2000). Zygotes from *unc-59(e1005)* often failed cortical rotation or rotated intermittently (Figure 2A, B, D and Supplemental Movie 1). Zygotes depleted of UNC-59 or from *unc-59(e1005)* worms were statistically indistinguishable with respect to maximal and average rotation velocities, and the duration and displacement of rotation (Figure 2D, F& G).

Septin function is thought to be fully removed by depleting one septin or mutating one of the two genes; heterotetramerization of UNC-59 and UNC-61 is thought to stabilize each protein (John et al., 2007; Nguyen et al., 2000; Nishihama et al., 2011). Nevertheless, since differences in loss-of-function phenotypes exist (Hall et al., 2008) we wished to compare the loss-of-function allele of *unc-59* to a null allele of the other *C. elegans* septin gene *unc-61 (e228)*, which contains a premature stop codon and in which no UNC-61 protein is detected by blotting with an antibody recognizing the N-terminus (Nguyen et al., 2000). Zygotes from the *unc-61* mutant strain were statistically indistinguishable from UNC-59 depleted and *unc-59(e1005)* zygotes in all our measures: maximum and average velocity, duration, and distance (Figure 2). Specifically, they had lower maximum and average speeds than control cells (Figure 2A, E -G). In sum, our findings demonstrated that three septin perturbations are essentially identical, and that septins are required for normal cortical rotation in *C. elegans* zygotes. Since the septins are not directly implicated in motor-driven force generation, these results suggest that additional factors are involved in translating molecular-or polymer-scale chirality to the cellular level.

### The formin CYK-1 is asymmetrically distributed in a septin-dependent manner

Two possible factors have been recently implicated in anaphase chiral rotation in the *C. elegans* zygote: anterior-posterior polarity and anterior-posterior myosin bundles (Pimpale et al., 2020; Singh et al., 2019). Thin film active fluid theory demonstrated that net chiral rotation (as occurs in the zygote in anaphase) is sensitive to lengthwise (anterior-posterior) cell polarity, such as the displacement of the division plane from the midplane of the cell, or anterior-posterior asymmetry of cortical contractility (Pimpale et al., 2020). The contribution of anterior-posterior polarity to cortical rotation is unlikely to explain rotation failure in septin loss-of-function cells, though, because septins are dispensable for zygote anterior-posterior polarity (Davies et al., 2016; Nguyen et al., 2000).

Another factor recently implicated in anaphase chiral rotation is the presence and integrity of “longitudinal” actomyosin bundles, extending between the anterior and posterior poles of the cell perpendicular to the cytokinetic ring (Singh et al., 2019). These bundles are assembled downstream of RhoA activity, but the source of F-actin within them was not examined. We hypothesized that the formin CYK-1, the only Dia family formin specific to cytokinesis in *C. elegans* (Davies et al., 2018), contributes to chiral cortical rotation. Since CYK-1 is dispensable for anterior-posterior polarity (Severson et al., 2002; Sönnichsen et al., 2005), perturbation of CYK-1 allows interrogation of F-actin-based structures without perturbing division plane placement.

We began by examining the distribution of CYK-1 using cells expressing CYK-1 fluorescently tagged at its endogenous locus (CYK-1::GFP; (Hirani et al., 2019). The invariant handedness of anaphase cortical rotation suggested that CYK-1 localization would be asymmetric, so we tested whether the distribution of CYK-1::GFP was polarized. CYK-1::GFP was slightly but consistently enriched in the zygote posterior immediately prior to anaphase onset (Figure 3A-C).

To test whether loss of septin function could perturb cortical rotation by affecting CYK-1, we compared CYK-1::GFP polarization in control cells to its localization in cells with incomplete septin function. We next depleted UNC-59 from cells expressing CYK-1::GFP to test the effect of loss of septin function on CYK-1 localization. Following UNC-59 depletion, the posterior enrichment of CYK-1::GFP was eliminated (Figure 3B, C). From this, we concluded that the septins are required for the polarization of CYK-1. By contrast, the cortical abundance and anterior enrichment of UNC-59::GFP was not affected by depletion of CYK-1 (Fig. S3 A).

Septins could impact the polarization of CYK-1 cortical enrichment indirectly, or more directly, by controlling its localization or stability. To define CYK-1 cortical dynamics, we first performed time-lapse imaging of CYK-1::GFP and measured fluorescence intensity on the cortex, correcting for the calculated loss of fluorescence due to photobleaching (Figure 3 supplement B-F). We found that leading up to anaphase onset and for the first minute of anaphase, when rotation occurs, levels of CYK-1::GFP on both the anterior and posterior poles of the cell increased over time (Figure 3 supplement, panel E). The relative enrichment of the formin on the posterior cortex also steadily rose (Figure 3D). By contrast, when UNC-59 was depleted, CYK-1::GFP remained uniformly distributed (Figure 3D). In sum, these results demonstrate that the formin CYK-1 is enriched in the posterior of the *C. elegans* zygote in a manner dependent on septins. This polarization suggested to us that CYK-1 also contributes to zygote chirality.

### The formin CYK-1 and septins make distinct contributions to cortical rotation

The importance of anterior-posterior myosin bundles for chiral cortical rotation (Singh et al., 2019) and our observation of asymmetric CYK-1::GFP localization suggested that this formin contributes to chiral cortical rotation. This hypothesis was also supported by the observation that elimination of long F-actin bundles via inhibition of formins blocked chiral cytoskeletal morphologies in adherent mammalian cultured cells (Tee et al., 2015). To test whether the formin CYK-1 is important for chiral contractility in the *C. elegans* zygote, we used RNAi to slightly deplete CYK-1 such that all cells still completed cytokinesis, and quantified cortical rotation. Like septin loss-of-function, CYK-1 depletion reduced the proportion of zygotes exhibiting normal rotation (Figure 4 A-C, G, H, and Figure 4 supplement). Unlike septin perturbations however, CYK-1 depletion did not reduce maximal or mean rotation velocity (Figure 4E-F), but notably, in many CYK-1 depleted embryos, cortical rotation occurred with opposite chirality or alternating chirality (Figure 4A-C and Supplemental Movie 1). Furthermore, both the duration of rotation and therefore total displacement were almost always (9/10) higher in CYK-1 depleted zygotes than in wild-type zygotes (Figure 4E-H). The wide range of abnormal cortical rotation effects following CYK-1 depletion likely reflects the incomplete and inconsistent cytoskeletal perturbations resulting from mild depletion of this essential protein. In sum, wild-type levels of the formin CYK-1 are dispensable for cortical rotation itself, but necessary for the consistent chirality of rotation.

The enrichment of the actomyosin cortical cytoskeleton in the anterior of the *C. elegans* zygote suggests a mechanism for the chirality of cortical rotation: an asymmetric accumulation of torsional stress (Pimpale et al., 2020). We hypothesized that CYK-1 depletion affects the handedness of cortical rotation by impacting cytoskeletal network polarization. We first tested whether CYK-1 depletion affects network polarization using the F-actin marker LifeAct and comparing LifeAct fluorescence in the anterior and posterior of the zygote in early anaphase. LifeAct fluorescence signal was indeed enriched in the anterior of control cells, and the average polarization of CYK-1 depleted cells was significantly lower (Fig. 4I). We next asked whether the subset of CYK-1 depleted cells that exhibited cortical rotation with continually or alternating reversed handedness had significantly lower F-actin polarization, and found this to be the case (Fig. 4I, filled black circles). These results support the hypothesis that the anterior-posterior polarity of the cortical cytoskeleton influences the chirality of cortical rotation in anaphase, possibly via impacting the accumulation of torsional stresses.

Our results above suggested that septins and CYK-1 contribute to chiral cortical rotation in distinct ways: septins help translate contractility into whole-cell chiral movement, and the levels and subcellular distribution of the formin CYK-1 contributes to the chirality of this contractility. Next, we tested whether septins and CYK-1 contribute in distinct ways by combining perturbations of septins and CYK-1. We depleted CYK-1 from *unc-61(e228)* zygotes and found that collectively, zygotes in which both CYK-1 and UNC-61 septin were perturbed exhibited more severe defects than those depleted of CYK-1 only; the average right-handed rotation velocity was essentially zero (Figure 4A, D, black line). This was because while some cells exhibited normal right-handed cortical rotation, others exhibited abnormal left-handed rotation. In general, rotation was slow, intermittent and often alternating between right-and left-handed directions (Figure 4A, D). When directionality information was discarded, and speed was simply compared, zygotes in which both CYK-1 and UNC-61 septin were perturbed were statistically indistinguishable from *unc-61(e228)* zygotes (Figure 4E, F). These results support the hypothesis that septins and the formin CYK-1 contribute in distinct fashions to chiral cortical rotation: the formin directs the handedness of rotation while septins translate cytoskeletal chirality to whole-cell cortical rotation.

## Discussion

Here, we show that the septins and a formin are required for chiral cortical rotation during anaphase in the *C. elegans* zygote. Specifically, rotation often fails when septins are depleted or mutated, and CYK-1 depletion did not eliminate cortical flows but altered their chirality. Together, this work advances our understanding of how molecular-scale chirality is scaled up to the level of cell-scale behaviors. Formins related to CYK-1 have been implicated in the chirality of adherent mammalian cells (Tee et al., 2015), but milder perturbations of formins will be necessary to discriminate between the requirement for formins for mammalian cells’ overall cytoskeletal abundance and network architecture versus its chirality. Septins may play a role in cellular chirality in other species, but since other animals have at least 5 septin genes and form combinatorial septin oligomers, achieving loss of septin function is difficult.

How does the cortex of the *C. elegans* zygote rotate inside of the eggshell? Our work connecting the septins and the formin CYK-1 to rotation, guided by published observations of cultured mammalian cells (Tee et al., 2015), suggests an explanation for this phenomenon (Figure 5). In mammalian cells, crosslinked perpendicular F-actin bundles accumulate the torsional stress of F-actin rotation as it is polymerized from membrane-anchored formins (Tee et al., 2015) (Figure 5A). Without membrane anchoring, the formin would freely rotate around the associated F-actin over the course of polymerization (Jégou et al., 2013; Mizuno et al., 2018; Zimmermann et al., 2017), and torsional stress would not accumulate. Accumulated stress is overcome by actomyosin contractility and relieved by chiral displacement of initially-radial bundles (Tee et al., 2015). Rotation speed in the *C. elegans* zygote is intermediate to that of NMMII movement and CYK-1 formin single molecule movement thought to be in concert with F-actin polymerization (Higashida et al., 2004; Li and Munro, 2020; Melli et al., 2018). This further supports the idea that chiral movement is not purely driven by NMMII motoring, but rather results from the release of torsional stresses in the network. The same conclusion is reached in a preprinted manuscript on the roles of CYK-1 in chiral cortical movements in *C. elegans* embryos (Middelkoop et al., 2021). When the dorsoventral axis of adherent mammalian cells is equated to the anterior-posterior axis of the *C. elegans* zygote, we would predict that formins would be enriched in the posterior; we indeed observe a slight but consistent posterior enrichment (Figure 3). We hypothesize that formins in the posterior generate F-actin bundles that emanate towards the cell anterior (Figure 5B, orange). The cytokinetic ring then comprises perpendicular bundles (Figure 5B, gold) running circumferentially and intersecting with the posterior-to-anterior bundles. Accumulated chiral torsional stress in the latter bundles and mechanical coupling between the two populations of F-actin would bias contractility to drive movement with right-handed chirality.

**Figure 5:**
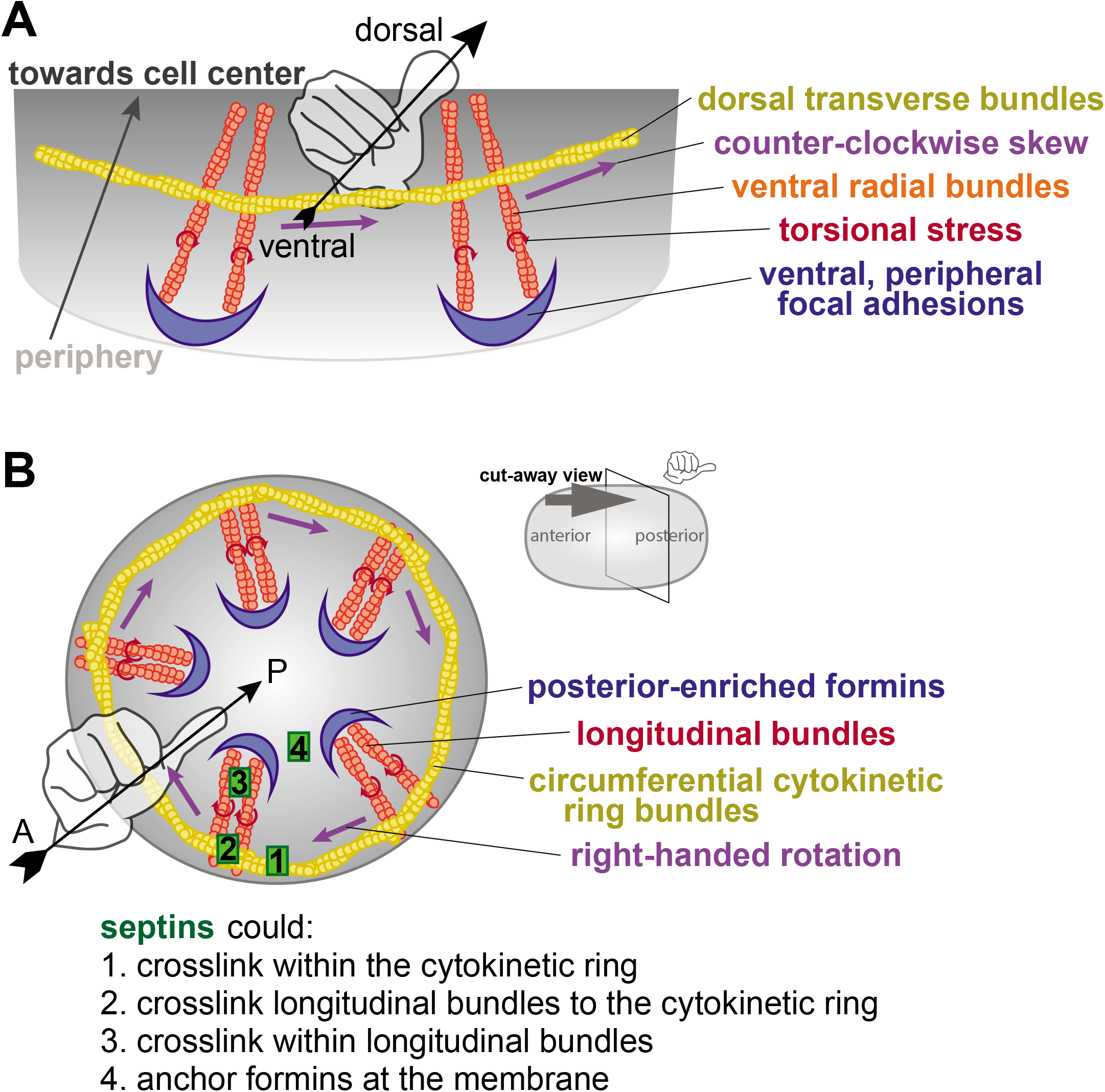
A model for the actin cytoskeleton structure in *C. elegans* zygotes. (A) Schematic of the hypothesized structure of the actin cytoskeleton that drives rotation in adherent mammalian cells (adapted from Tee et al., 2015). Peripheral, anchored formins (blue) generate radial F-actin (orange) under torque (red) and coupled to circumferential bundles (yellow) whose contractility leads to a counter-clockwise skew (purple). (B) Cut-away perspective view into the posterior pole of the *C. elegans* zygote illustrating hypothetical mechanism for cortical rotation: Membrane-anchored, posterior-enriched formins (blue) generate longitudinal (posterior-to-anterior) bundles that interact with circumferential cytokinetic ring actin bundles (yellow). Contraction in the ring could relieve chiral torque (red) within posterior-to-anterior bundles (orange), generating a right-handed rotation (purple). Septins (green) could contribute to this phenomenon by localizing/anchoring formins to the posterior focal adhesions or crosslinking within or between actin bundle populations.

How do septins translate non-muscle myosin II-driven contractility into cortical rotation? Septins are not directly implicated in the formation or translocation of F-actin filaments and bundles, but rather in network connectivity. Septins crosslink the cortical cytoskeleton via interactions with F-actin, NMMII, and anillin (reviewed by Spiliotis, 2010); this is expected to cause the accumulation of stresses in the network (Figure 5B, green 1-3). Furthermore, septins could anchor formins at the plasma membrane (Figure 5B, green 4), limiting formin rotation around F-actin during polymerization, and thus leading to the accumulation of torsional stresses within the network (Jégou et al., 2013; Mizuno et al., 2018; Tee et al., 2015; Zimmermann et al., 2017). We predict that reduced connectivity in any or all of these aspects of the network reduces the accumulation of polymerization-associated torsional stress so that cytokinetic contractility does not lead to a release of tension and thus cortical rotation.

How does the formin CYK-1 contribute to the chirality of cortical flow and how does rotation occur with reversed (left-handed) chirality? The chirality of F-actin is invariant, as is the chirality of the torsional stress generated by formin-driven polymerization of actin into a highly crosslinked network. However, CYK-1 is only slightly enriched in the posterior, and mild depletion of CYK-1 may randomize or eliminate this asymmetry. Depletion of CYK-1 itself diminishes anterior enrichment of F-actin, possibly reflecting an alteration of the balance of formin versus Arp2/3-based F-actin in the two poles of the cell (Rotty and Bear, 2014). Preferential or even intermittent interaction between circumferential cytokinetic ring F-actin and longitudinal bundles extending from the anterior towards the posterior would result in left-handed chirality of the anaphase contractile network. We thus predict that when CYK-1 polarization is eliminated or reversed, cortical rotation handedness will be reversed. Indeed, when upstream regulation of anterior-posterior polarity is perturbed via depletion of PAR proteins or Cdc-42, cortical rotation with reversed handedness is exhibited by some embryos (Schonegg et al., 2014).

Chiral cortical rotation in the zygote represents the first appearance of handedness in a cell with radial symmetry around a single axis. Failure to establish body axes leads to early lethality (Belmont et al., 2004; Ware et al., 2004). Exploration of whether zygote chiral rotation is essential for viability has been limited because all other proteins reported to be required for this chirality are central to basic cell biological processes (*e*.*g*. tubulin, centrosome maturation machinery, the RhoGEF ECT-2, and non-muscle myosin II heavy chain; (Schonegg et al., 2014)). Although strains bearing septin loss-of-function alleles are viable, they exhibit a broad range of phenotypes including arrested development, uncoordinated movement, and vulval protrusion indicating tissue morphogenesis defects (Nguyen et al., 2000). It will be interesting to test whether the severity of cortical rotation defects following septin loss-of-function correlates with decreased viability or randomized or reversed body axis handedness, which is tolerated in *C. elegans* as it is in humans (Goldstein and Hird, 1996; Kosaki and Casey, 1998; Wood, 1991).

## Supporting information

Zaatri et al Supplemental Movie 1

Zaatri et al Supplemental Movie 2

## Acknowledgements

The authors are grateful to Julie Canman and her lab for characterizing and providing septin mutant strains. We thank Vincent Boudreau for creating tools in ImageJ, Iris Brammer for assistance in data collection, and Dylan Ray for technical assistance. For valuable discussion, we thank Bill Wood, Stephan Grill and Teije Middelkoop, and all the members of the Maddox labs. We thank Richard Cheney, Daniel Cortes, Bob Goldstein, Michael Werner, and Ben Woods for critical reading of this manuscript.

## Figure Legends

**Supplement to Figure 2:**
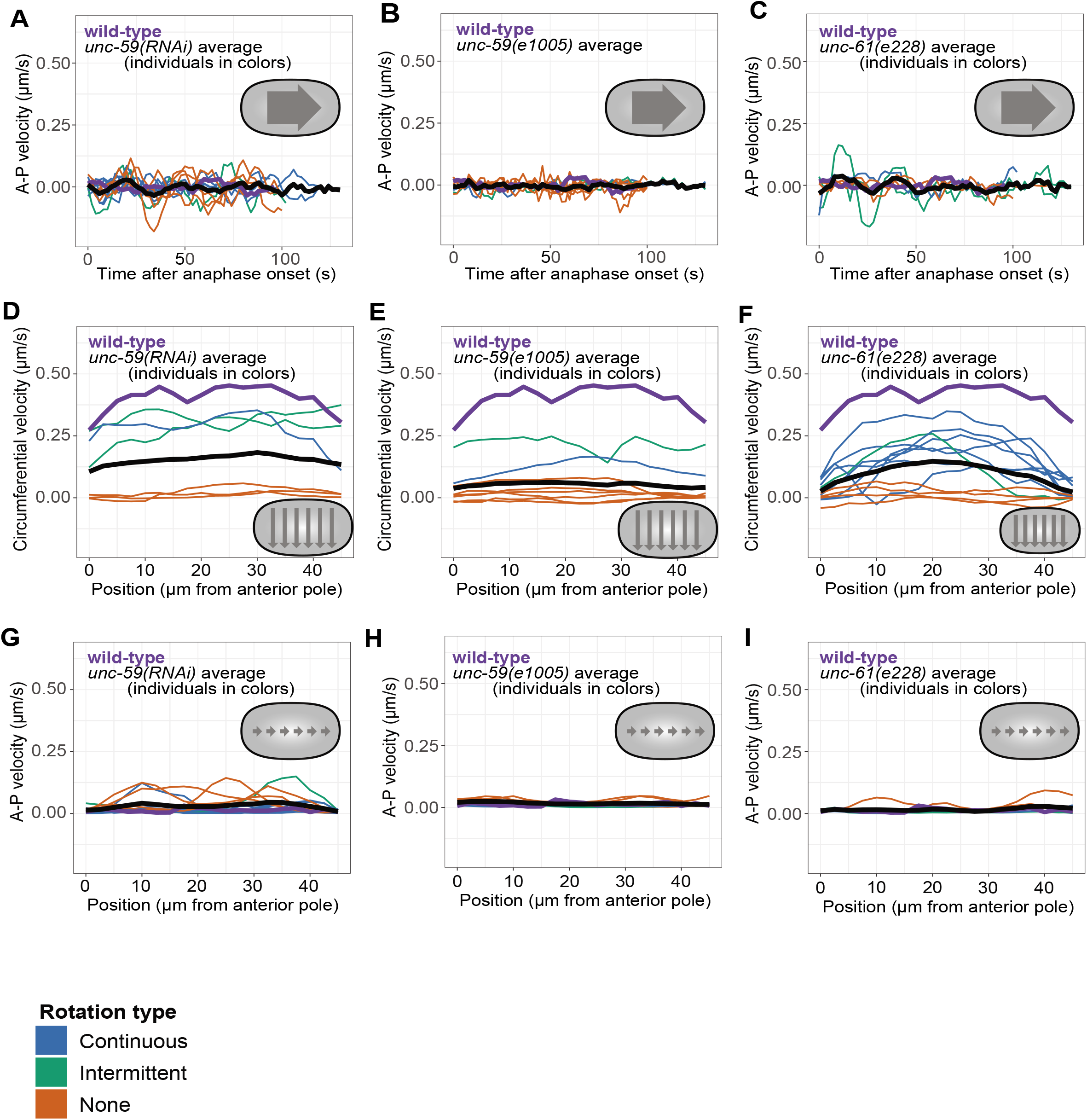
Anterior-to-Posterior and binned velocities of septin loss of function cells. (A-C) Mean and variance (+/− one standard deviation) of circumferential velocities over time averaged across the anterior-posterior axis. (D-I) Posterior-directed velocities over time for *unc-59(RNAi), unc-59(e1005)*, and *unc-61(e228)* cells (n = 12, n = 9, n = 6, respectively). Gold, green, blue and dashed colored lines indicate individuals of the corresponding color-coded phenotype; black: septin perturbation population average; purple: control average. (G-I) Mean and variance (+/− one standard deviation) of posterior-directed velocities over time averaged across the top-bottom axis. (J-L) Binned circumferential velocities. Populations and color denotations are the same as panels D-F. (M-O) Binned posterior-directed velocities. Populations and color denotations are the same as panels D-F.

**Supplement to Figure 3:**
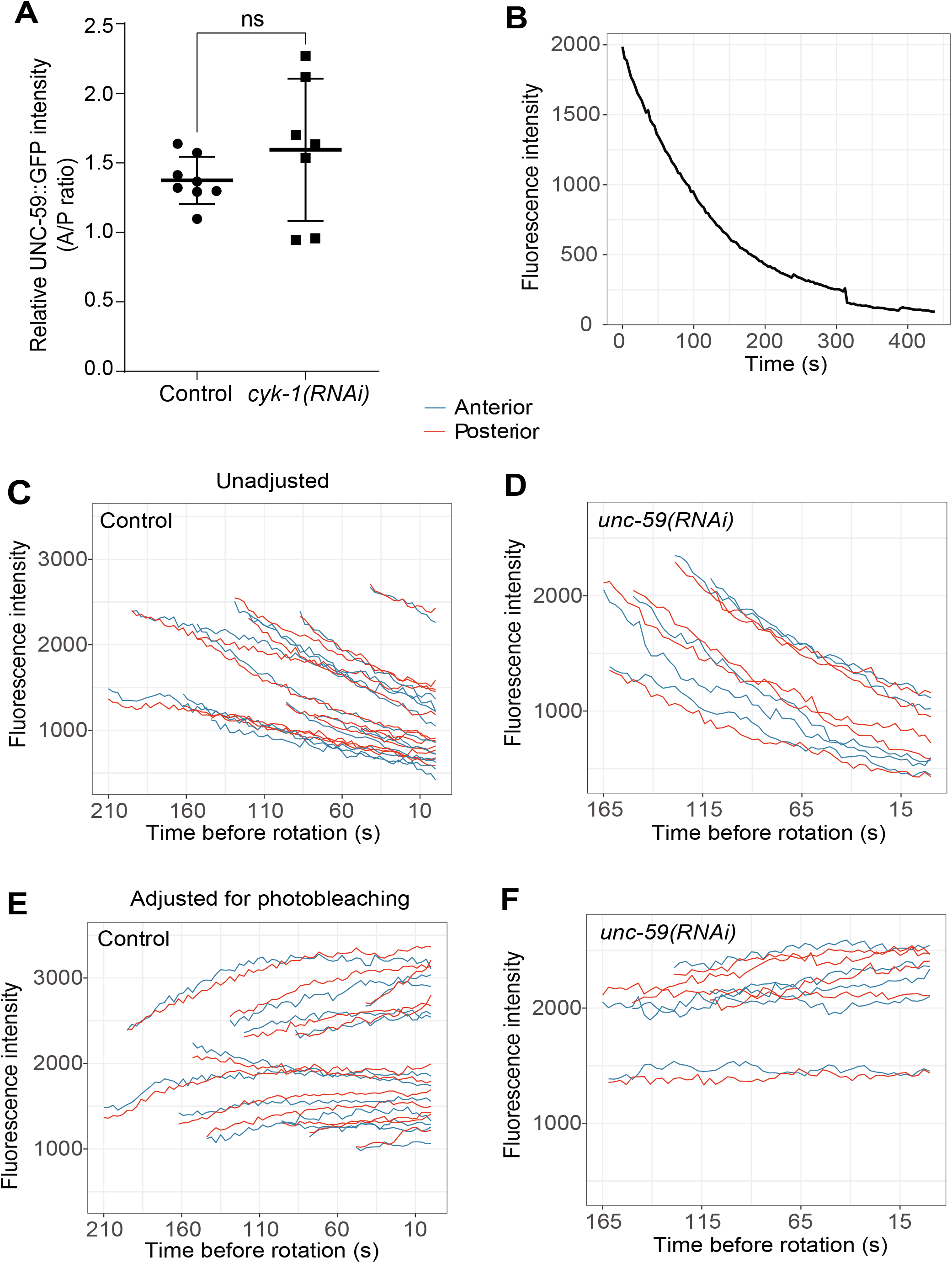
Controls for Figure 3. (A) Mean intensity ratio (A/P) of UNC-59::GFP just prior to anaphase onset for control and *cyk-1(RNAi)* cells. Dots are individuals; horizontal bar is population mean; vertical notches are 95% confidence interval; ns = not significantly different. (B-F) Photobleaching adjustment for CYK-1::GFP cells. (B) Averaged fluorescence intensity CYK-1::GFP in cells treated with azide (n=10) (B-C) Cortical fluorescence intensities over time in CYK-1::GFP control (C, n =12) and UNC-59 depleted (D, n = 5) cells. (E-F) Photobleaching-corrected CYK-1::GFP intensities for the same cells as in B and C. Movies were recorded on the same settings as those made to build the decay model in (B); the decay coefficient was used to calculate and add in the intensity that was lost to photobleaching.

**Supplement to Figure 4:**
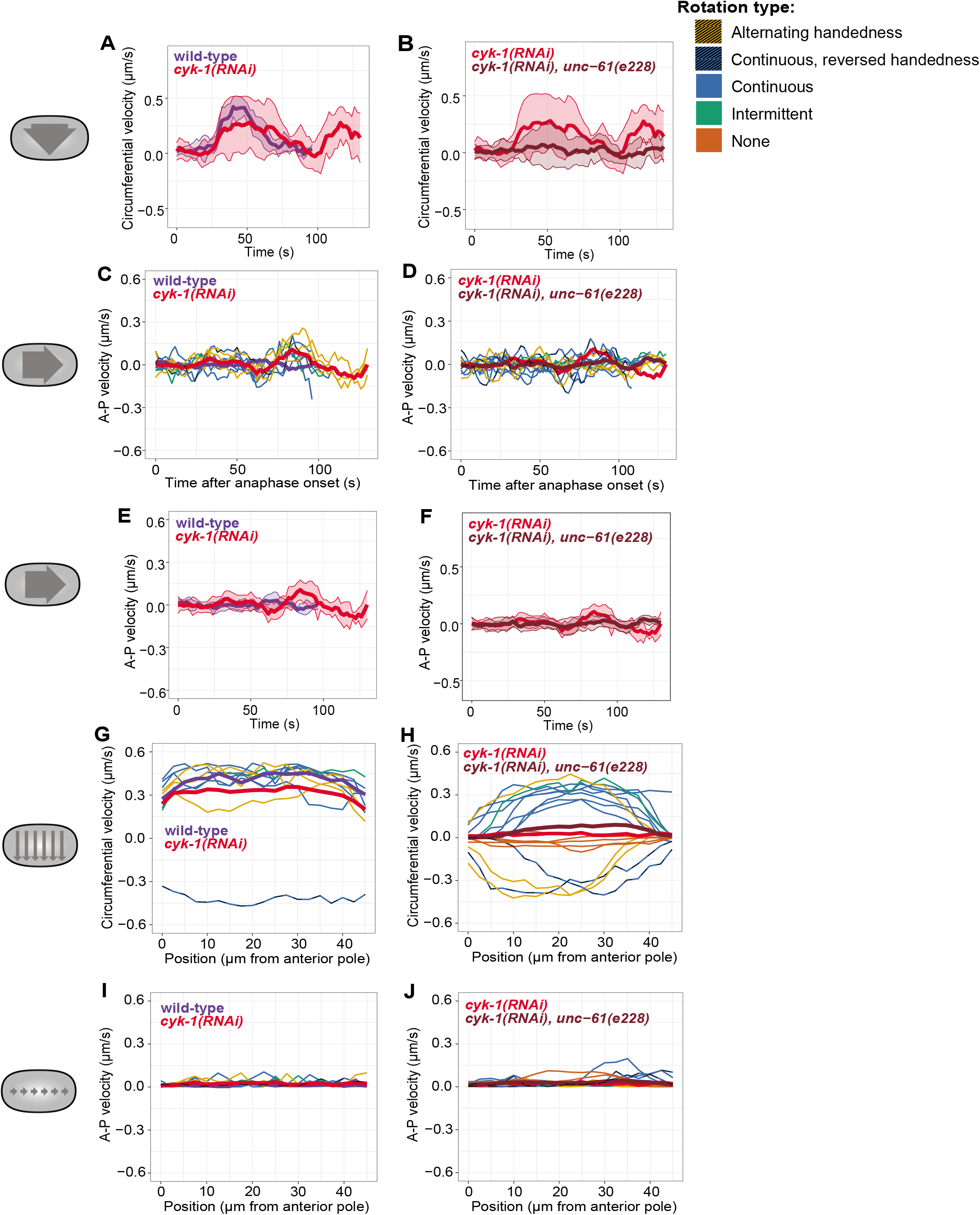
AP and binned velocities for *CYK-1* loss of function cells. (A and B) Mean and variance (+/− one standard deviation) of circumferential velocities over time averaged across the anterior-posterior axis. (C-F) Posterior-directed velocities over time for *cyk-1(RNAi)* and (*cyk-1(RNAi)*; *unc-61*(e228)) populations (n = 10, n = 18). Brown: (*cyk-1(RNAi)*; *unc-61(e228)*) population average; purple: wild-type average; red: *cyk-1(RNAi)* average. (C and D) Gold, green, blue and dashed colored lines indicate individuals of the corresponding color-coded phenotype. (E and F) Mean and variance (+/− one standard deviation) of posterior-directed velocities over time averaged across the top-bottom axis. (G and H) Binned circumferential velocities. Populations and color denotations are the same as panels A-B. Dashed line indicates reversed handedness. (I and J) Binned posterior-directed velocities. Populations and color denotations are the same as panels A-B.

**Supplemental Movie 1**

Time lapse of cells exhibiting (A) continuous cortical rotation (wild-type), (B) intermittent cortical rotation (*unc-59(e1005)*), (C) no cortical rotation (*unc-59(e1005)*), (D) cortical rotation with reversed handedness (*cyk-1(RNAi)*), and (E) alternating cortical rotation (*cyk-1(RNAi)*). Images were acquired at 2 second intervals; playback is 12 x realtime. Scale bar = 10 µm.

**Supplemental Movie 2**

Time lapse of cells exhibiting (A) continuous cortical rotation (wild-type), (B) intermittent cortical rotation (*cyk-1(RNAi); unc-59(e1005)*) and (C) cortical rotation with reversed handedness (*cyk-1(RNAi); unc-59(e1005)*). Images were acquired at 2 second intervals; playback is 12 x realtime. Scale bar = 10 µm.

